# *Transplant-Agents*: A Multi-Agent Artificial Intelligence Framework for Reproducibility Assessment of Post-Transplant Risk Prediction and Rejection Biomarkers

**DOI:** 10.1101/2025.07.10.664265

**Authors:** Sirui Ding, Sanchita Bhattacharya, Minnie M. Sarwal, Marina Sirota, Atul J. Butte

**Affiliations:** Bakar Computational Health Sciences Institute, University of California, San Francisco, CA, USA; Division of Transplant Surgery, Department of Surgery, University of California San Francisco, San Francisco, California, USA

## Abstract

Reproducible biomarker identification and transplant rejection risk prediction remain fundamental yet unsolved challenges in transplantation medicine. Traditional approaches rely on hypothesis-driven analyses and domain expertise, limiting scalability and generalizability across diverse populations. We introduce *Transplant-Agents*, a data-driven multi-agent AI framework integrating large language models (LLMs) with machine learning algorithms for automated biomarker identification and rejection risk prediction. Agents interact through structured, iterative dialogue governed by predefined rules and criteria, converging on optimal biomarker sets reproducible across multiple iterations. We evaluated three multicenter clinical trial transplant datasets from ImmPort, comprising 683 patients across kidney, liver, and heart transplant cohorts. *Transplant-Agents* achieve AUROC scores of 0.93, 0.88, and 0.88. Feature importance analysis further confirms the stability, interpretability and potential generalizability of identified biomarkers. This work demonstrates that AI-agent frameworks can reliably reproduce established transplant biomarkers while enabling transparent, validated, and standardized risk prediction pipelines.

## Introduction

The integration of artificial intelligence into clinical decision-making has transformative potential, yet translating predictive models into interpretable, actionable insights remains a fundamental challenge in biomedical AI. This challenge is particularly acute in organ transplantation, where risk stratification directly impacts patient outcomes and resource allocation, and where clinicians require not only accurate predictions but also mechanistic understanding of the underlying risk factors for clinical decision support^1^. For instance, identifying non-invasive rejection biomarkers from blood and urine to predict rejection is an essential topic for the transplant community, from both clinical and research perspectives. Though transplant clinicians usually depend on the graft biopsy ^2^ to identify rejection risk after transplant^3^, there is an unmet need to identify rejection through non-invasive monitoring and additionally to utilize the biomarkers from these biofluids to predict rejection in the organ, prior to graft dysfunction. It is also important to identify biomarkers with both high sensitivity and specificity, so that rejection diagnosis is not confounded by additional factors that can influence graft function, such as the quality of the organ, delayed graft function, demographic variables, and recipient co-morbidities, sensitization, waiting list time^4–6^, and Model for End-Stage Liver Disease (MELD) scores^7,8^ and residual kidney function^9^. Many current methods for diagnosing graft rejection have significant limitations. Subclinical rejection often goes undetected, and molecular rejection can be present even in the absence of histological injury^10^, making it difficult to assess and predict the evolution of graft injury over time. Fixed scoring systems lack adaptability and generalizability, failing to accommodate shifts in patient cohort distributions driven by demographics, socioeconomic status, familial factors, and complex disease patterns. Conventional approaches that restrict biomarker analysis to a predefined set of input variables are further constrained in their ability to identify novel biomarkers and risk factors.

The increasing accessibility to publicly available clinical trials and research data^11^ including patient-level demographics, multi-omics datasets, and clinical laboratory measurements now offers an opportunity to validate the output from the traditional calculators into data-driven, adaptive systems. ML, deep learning (DL) and other AI-based biomarker identification methods have been applied across domains such as oncology and other disease areas ^12,13^ but remain to be broadly applied to transplant research.

To strengthen the reproducibility and rigor of expert-designed clinical biomarker discovery and conventional AI models in transplantation, we developed *Transplant-Agents*, a data-driven multi-agent AI framework for rapid and robust biomarker identification and risk prediction. The system comprises of two collaborating agents: an ML Engineer Agent, which calls ML model APIs to optimize biomarker features, and a Domain Expert Agent, which applies domain-specific knowledge and constraints to ensure biomedical relevance and interpretability. The agents interact through structured, iterative dialogue governed by predefined rules and criteria, converging optimal biomarker sets that remain reproducible across multiple iterations.

We leveraged the studies from ImmPort^8,14^ publicly accessible immunology data portal that contains subject-level biomarker data and corresponding clinical phenotypes from the transplantation domain. For prediction, the agent selected and implemented a tree-based model, Light Gradient Boost Model (LightGBM^15^), as the predictive tool for its accurate performance and interpretability on structured clinical data. Post-hoc analysis provides feature importance and biomarker interaction insights, ensuring transparency. We benchmarked Transplant-Agents against expert-designed clinical risk calculators, conventional feature selection methods, and baseline machine learning models across all three transplant predictive tasks.

This study presents three key advancements: First, we developed a multi-agent AI system, *Transplant-Agents*, powered by LLMs, to automatically identify biomarkers and predict risk for organ transplant. Second, through transplant clinical research datasets, *Transplant-Agents* train models which are compared against expert-designed rules, e.g., MELD score and baseline machine learning models for predicting transplant rejection. The system offers a robust and interpretable AI framework suitable for clinical use. Third, *Transplant-Agents* reproducibly identified previously validated biomarkers from the literature, demonstrating the capacity for consistently corroborating published biomarker associations and provide rationale for the potential to *Transplant-Agents* to support new scientific discoveries.

## Methods

### Data pre-processing

We used longitudinal data from three CTOT consortium studies—CTOT-01 (kidney), CTOT-14 (liver), and CTOT-05 (heart)—comprising 683 patients across U.S. and Canadian sites, obtained from the ImmPort portal^14^. Each study included static variables (demographics, medical history) and longitudinal variables (clinical and molecular measurements), with post-transplant rejection as the primary prediction endpoint. Only longitudinal measurements collected before the rejection event were used as inputs, summarized into minimum, maximum, and most recent values per time series. These summary features, combined with the static variables, were used to predict rejection and identify associated salient variables. Variables with over 30% missing data were excluded; remaining missing entries were imputed with zeros.

### Framework Overview

In typical biomedical scenarios, including experiment and clinical trials, data collection typically comprises a combination of static and longitudinal data. Static data usually comes from demographic information, medical history, and baseline clinical information. Longitudinal data such as laboratory tests and transplant-specific clinical tests are collected at multiple time-points during pateint follow-up at pre-and post-transplantation. We employed a simple yet effective pre-processing feature engineering method to derive the maximum, minimum, and latest values from longitudinal variables and concatenated them with static variables^16^. The processed variables serve as input into the *Transplant-Agents* framework as shown in Figure 1(a), which has two specialized agents on ML and domain expertise. The ML agent served as an ML engineer, and the domain expertise agent will serve as a domain expert, e.g., transplant clinician. The computational tools are available for ML agents, for example, LightGBM in our design and implementation, and the feature importance calculator, which calculates feature importance from the tree-based model (e.g., LightGBM). The ML agent calls the ML model to predict the risk of the primary target, e.g., transplant rejection. Then the feature importance calculator is called to rank the features. Through prompt learning, the ML engineer agent will make the decision to add or delete features to improve prediction performance based on the feature importance (determined by the cumulative gain) and predictive results. Then, the domain expert agent will review the add or delete operation to make sure it follows the basic biomedicine and clinical principles and knowledge. Multiple rounds of discussion like this will be held between the ML engineer and domain expert agents until a consensus is reached. The output of *Transplant-Agents* identified important biomarkers and predicted rejection risk.

**Figure 1.**
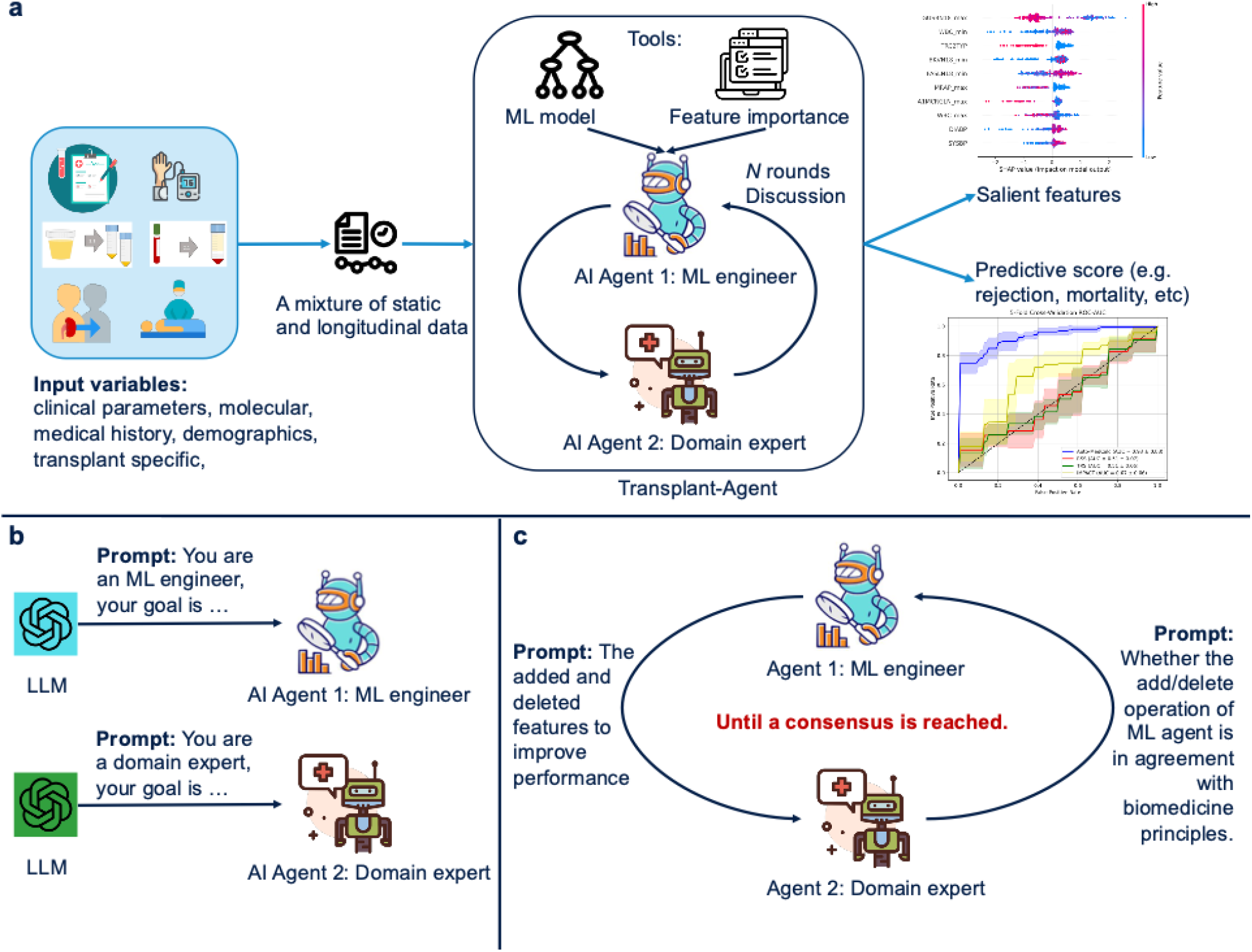
Overview of the proposed Transplant-Agents. **a:** Workflow of the automated biomarker identification and risk score prediction with a multi-agent AI system. The input is a mixture of static and longitudinal data. The output is salient features and a predictive score. **b:** Building a specialized AI agent from a large language model (LLM) with prompt learning. **c:** Collaboration between agents is achieved by multiple rounds of discussion until a consensus is reached.

**Figure 2.**
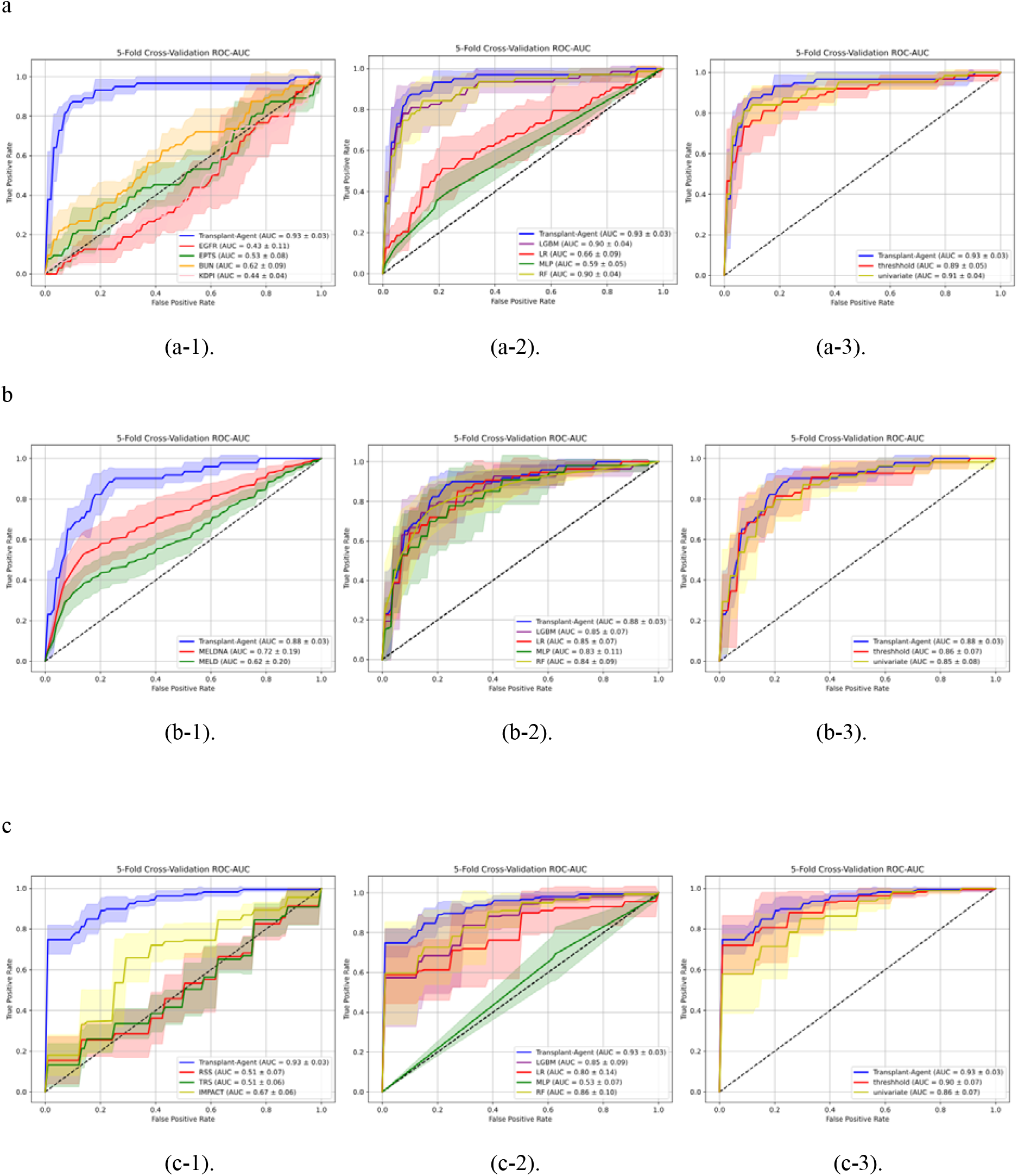
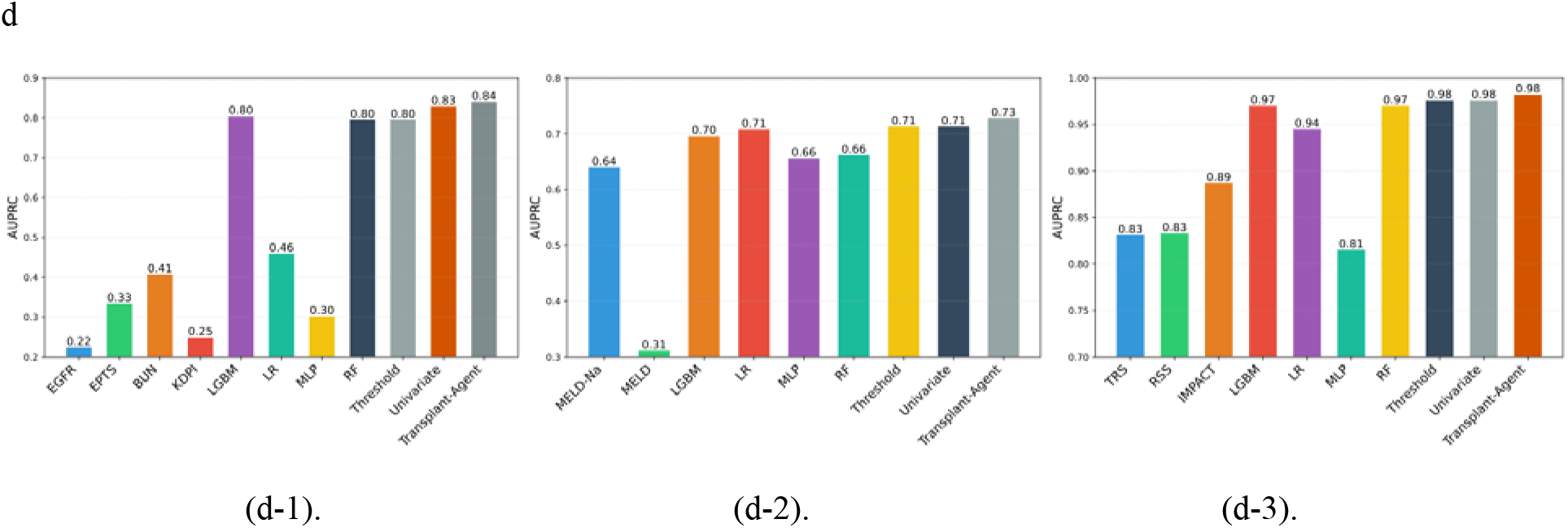
Performance compared with human-designed rules, baseline ML models, and feature selection methods, respectively. **a.** AUROC comparison on kidney transplant rejection prediction. a-1, a-2, a-3 indicate the performance compared with human-designed baselines, ML baselines and feature selection baselines respectively. **b.** AUROC comparison on liver transplant rejection prediction b-1, b-2, b-3 indicate the performance compared with human-designed baselines, ML baselines and feature selection baselines respectively. **c**. AUROC comparison on heart transplant rejection prediction c-1, c-2, c-3 indicate the performance compared with human-designed baselines, ML baselines and feature selection baselines respectively. **d**. AUPRC results comparison across kidney, liver, and heart rejection prediction. d-1, d-2, d-3 indicate the performance compared with baselines on kidney, liver, and heart respectively.

### Building agents from LLM

The AI agents are built from the LLM with prompt learning as shown in Figure 1(b) and Figure 4. We designed a ‘Role-Expertise-Task’ prompt template to build the specialized agent^17^. The ‘Role’ defines the character the agent will act in the task, e.g., ML engineer, domain expert, clinician, etc. The ‘Expertise’ specifies the skills or scientific expertise the agent will have, e.g., feature selection, model building, kidney transplant, etc. The ‘Task’ indicates the goal of the agent, or the task the agent should complete, e.g., diagnose disease (prediction/classification task), adjust the input variable set to optimize diagnosis performance (feature selection task), analyze features from a clinical perspective (domain critic task), etc.

For the ML engineer agent, we assign the ‘Role’ as ML engineer. The ‘Expertise’ of an ML engineer agent is building a machine learning pipeline and feature engineering, which are common ML skills. And the ‘Task’ is to diagnose the disease, optimize the diagnosis performance, and provide an interpretation of the results with XAI techniques.

For the domain expert agent, we assign the ‘Role’ as the domain expert. The ‘Expertise’ of a domain expert can be kidney transplant, liver transplant, or heart transplant, etc. And the ‘Task’ is to check whether the input features are consistent with basic biomedicine knowledge and provide critical feedback. The details of the prompts we use are presented in Figure S1 of the supplementary file.

### Processing input data with agent

The input data consists of lab tests, biomarkers, demographic informatics, etc., which are categorical or numerical variables. Instead of directly using an LLM-driven agent to process the raw data, including both variable names and values, we design a paradigm to separately intake names and values into the agent, as shown in Figure 4(a). The agent will only process the input variable names, which can leverage the strength of LLM to understand the semantic and clinical meanings of each variable. The agent will use ML tools to directly intake the variable values and evaluate the performance of the current feature set on the designated task. ML tools can be some ML models, e.g., LGBM, etc. The disentangling of variable names and values eases the burden of the agent, which can just use tools, do analysis like a ‘brain’. After the ML tool is called to process the input variables, the agent will intake and analyze the results and the feature importance from the ML model.

### Tools usage for task completion

For the diagnosis task, the agent utilizes ML models as the core predictive tool. These models are trained on labeled datasets to learn complex patterns and relationships between input features and target outcomes. Commonly used models may include decision trees, random forests, support vector machines, gradient boosting algorithms (e.g., XGBoost, LightGBM), or DL architectures, depending on the nature and complexity of the problem.

For the feature analysis task, the agent employs feature importance tools to evaluate the contribution of each feature to the model’s prediction performance. This includes both filter-based methods (e.g., mutual information, correlation analysis), wrapper methods (e.g., recursive feature elimination), and model-based approaches. A typical tool for feature importance analysis is built-in feature importance from tree-based models, which quantitatively ranks features based on their impact on model accuracy or loss.

For the interpretability and visualization task, the agent adopts SHAP (Shapley Additive explanations) value-based methods. The SHAP package provides a unified framework to explain individual predictions by attributing them to input features in a theoretically grounded way, based on cooperative game theory. This enables both global and local interpretability, allowing users to understand not only which features are important in general, but also how each feature contributes to specific predictions.

### Collaboration between agents

The collaboration between the two agents—namely, the ML engineer agent and the domain expert agent—is designed as an iterative discussion process that facilitates knowledge exchange and decision consensus. These agents engage in a dynamic dialogue where each contributes to its specialized expertise to refine the overall analysis and conclusions.

Specifically, the ML engineer agent focuses on the technical aspects of model development, including feature engineering, model selection, and performance evaluation. Meanwhile, the domain expert agent provides critical contextual insights grounded in domain-specific knowledge, helping to validate the relevance and plausibility of model findings and feature interpretations.

During the collaboration, the ML engineer agent presents model outputs, such as diagnostic predictions and selected features based on importance rankings. The domain expert agent then reviews these outputs, questioning or endorsing specific features or model behaviors based on domain understanding. If discrepancies or uncertainties arise, the agents exchange feedback iteratively, prompting model adjustments or re-examination of the data.

This back-and-forth continues until both agents reach a consensus, harmonizing the technical rigor of machine learning with the nuanced understanding of domain expertise. This consensus-driven approach ensures that the final diagnostic decisions or feature interpretations are both statistically sound and practically meaningful.

As illustrated in Figure 1(c), this discussion method enhances the reliability and interpretability of the system’s outputs by leveraging complementary strengths of the two agents through cooperative interaction. Such a collaborative framework not only improves decision quality but also fosters transparency and trustworthiness in the deployment of AI-driven solutions.

### Implementation for Transplant Rejection

The *Transplant-Agents* framework is leveraged for scalability and modularity, allowing it to adapt to biomarker identification and diagnostic tasks by reconfiguring its agentic goals and toolsets. We validated our approach using organ transplantation—a domain with an acute clinical demand for precise risk stratification and non-invasive monitoring. Currently, clinicians and researchers prioritize the discovery of non-invasive biomarkers to preemptively detect graft rejection and optimize post-transplant outcomes. *Transplant-Agents* extend these clinical physician capabilities by autonomously distilling high-relevance biomarkers, architecting specialized diagnostic pipelines, and providing clinically grounded interpretations of the results.

To evaluate the framework’s ability to reproduce and identify clinical biomarkers, a primary objective was to predict transplant rejection by identifying critical non-invasive biomarkers from multi-modal inputs, including urine, blood, and vital signs, prior to the clinical onset of rejection. This aligns with the study goal of the original CTOT trials: transitioning from invasive biopsies to reliable, non-invasive molecular and physiological diagnostics. *Transplant-Agents* achieve this complex objective through a structured four-stage workflow, ensuring the fidelity of the identified biomarkers relative to original clinical trial findings.

### Building ML engineer agent

First, we build the ML engineer agent (Box 1), which plays a pivotal role in constructing a robust machine learning pipeline tailored for diagnosing post-transplant rejection. This agent, designed with expertise in developing diagnostic ML pipelines, leverages structured input features to ensure precise and reliable outcomes. For Task 1, the agent is responsible for training and testing a machine learning model using patient lab tests and supplementary clinical data. This process involves preprocessing the data, selecting appropriate algorithms, and evaluating model performance to accurately identify instances of post-transplant rejection. For Task 2, the agent focuses on optimizing the input variables by employing computational techniques to identify and prioritize high-relevance biomarkers. This optimization enhances the model’s diagnostic accuracy by reducing noise and focusing on the most impactful features, ultimately improving the reliability of the diagnostic process.

- **Role**: ML engineer.
- **Expertise**: Building diagnosis ML pipeline with structured input features
- **Task 1**: Given the patient lab tests and other information, train and test the machine learning model to diagnose post-transplant rejection.
- **Task 2**: Optimize the input variables to select the high-relevant biomarkers from computational perspective, which could further improve the diagnosis accuracy.

Box 1.

### Building transplant expert agent

Second, we build the transplant expert agent, which provides clinical feedback for the ML engineer agent. The transplant expert agent is designed with specialized expertise in organ transplantation, focusing on kidney, liver, and heart procedures, and possesses an in-depth understanding of the intricate relationship between biomarkers and post-transplant rejection. This agent acts as a critic from a clinical perspective, tasked with reviewing the variables selected by the ML engineer agent to ensure their relevance and scientific validity in the context of transplantation outcomes.

- **Role**: Kidney/Liver/Heart transplant expert
- **Expertise**: Expertise in organ transplantation, particularly understanding the relationship between biomarkers and post-transplant rejection.
- **Task 1**: Serve as a critic agent from the clinical aspect, review the selected variables by the ML engineer agent and provide scientific feedback with domain knowledge.
- **Task 2**: Clinically analyze the important features from the interpretable results. Identify the essential/novel biomarkers related with the post-transplant rejection.

Box 2.

### ML engineer and transplant expert discussions

The ML engineer agent and the transplant expert agent collaborate through multiple rounds of discussions, fostering an iterative process where the ML engineer proposes selected biomarkers, and the transplant expert evaluates these choices based on clinical significance, biological plausibility, and evidence from transplant immunology

For instance, the transplant expert agent may critique the inclusion of certain biomarkers, such as cytokine levels or donor-specific antibodies, by assessing their established roles in acute or chronic rejection, their temporal expression patterns post-transplant, and their potential as confounders in predictive algorithms. This collaborative dialogue ensures that the chosen variables align with clinical realities, enhancing the model’s applicability in real-world transplant scenarios and improving its ability to predict rejection risks accurately. Through this dynamic exchange, the transplant expert agent bridges the gap between data-driven insights and clinical expertise, ultimately refining the machine learning approach to better serve patient outcomes in organ transplantation.

### Tool Selection and Implementation

#### LGBM is chosen for diagnosis

The ML engineer agent selects and implements the ML model used for rejection diagnosis. As shown in Figure B1, the ML engineer agent selects the Light Gradient Boosting Machine (LGBM) as the predictive model based on the input data type, task type, and its knowledge as an ML engineer. Also, the ML engineer agent is asked to generate the code of training and testing LGBM as shown in Figure B2.

#### Feature importance from LGBM

Feature importance from LightGBM, calculated using metrics like gain or split, quantifies each feature’s contribution to the model’s predictions. The ML engineer agent uses these scores to optimize the feature set by prioritizing impactful features and reducing less relevant ones, improving model accuracy and efficiency. The agent also generates implementation code to extract and visualize feature importance using LightGBM’s feature_importance() method, integrating libraries like pandas and matplotlib for clear, reproducible results in the machine learning pipeline.

#### SHAP value analysis and visualization

SHAP (SHapley Additive exPlanations) value analysis is employed to interpret and analyze the results of a machine learning model by quantifying the contribution of each feature to individual predictions. Additionally, the ML engineer agent is tasked with generating implementation code to compute SHAP values using libraries like shap in Python, integrating with the model to calculate explanations and produce visualizations with tools like matplotlib or plotly.

#### *Transplant-Agents* is an Interpretable AI Agent System

*Transplant-Agents* is a white-box system, for which we can understand each action/decision the agents made. In the scenario of biomarker identification and disease diagnosis, the most essential action of the agent is to select highly relevant biomarkers for precise diagnosis. *Transplant-Agents* provides interpretation for each biomarker selection decision made by the ML engineer agent and the clinical analysis from the domain expert agent. As shown in Figure 3 (d), this is the screenshot of a one-round discussion between two agents. We can observe that the ML engineer agent returns the selected biomarkers, which can improve the diagnosis precision from the computational perspective. The transplant expert agent analyzed these biomarkers from a clinical standpoint to ensure they correspond to the transplant expertise principles. After the diagnosis made by the ML engineer agent, the domain expert agent can also analyze the results from a clinical perspective and identify potential new biomarkers, as shown in Figure 5.

**Figure 3.**
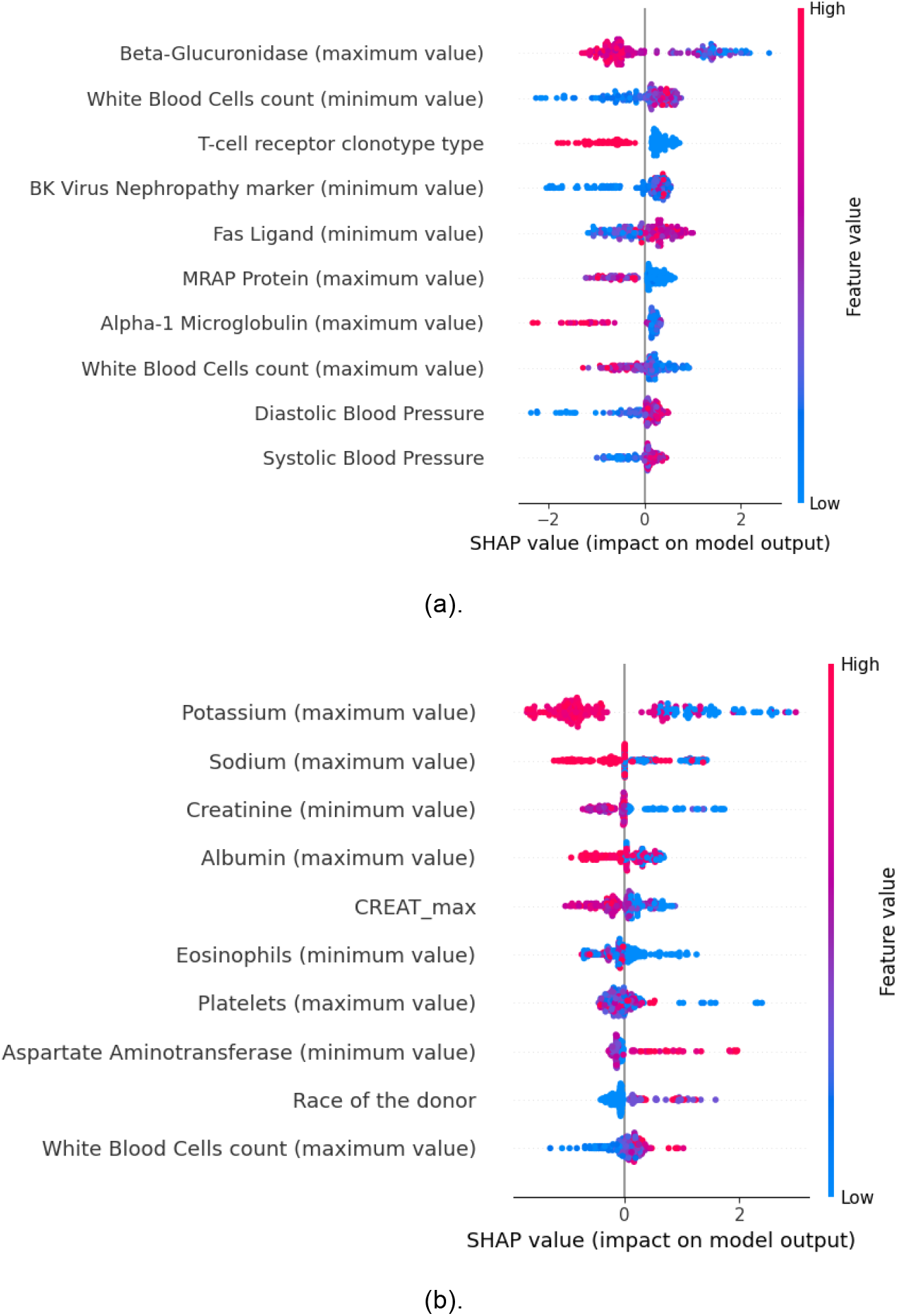
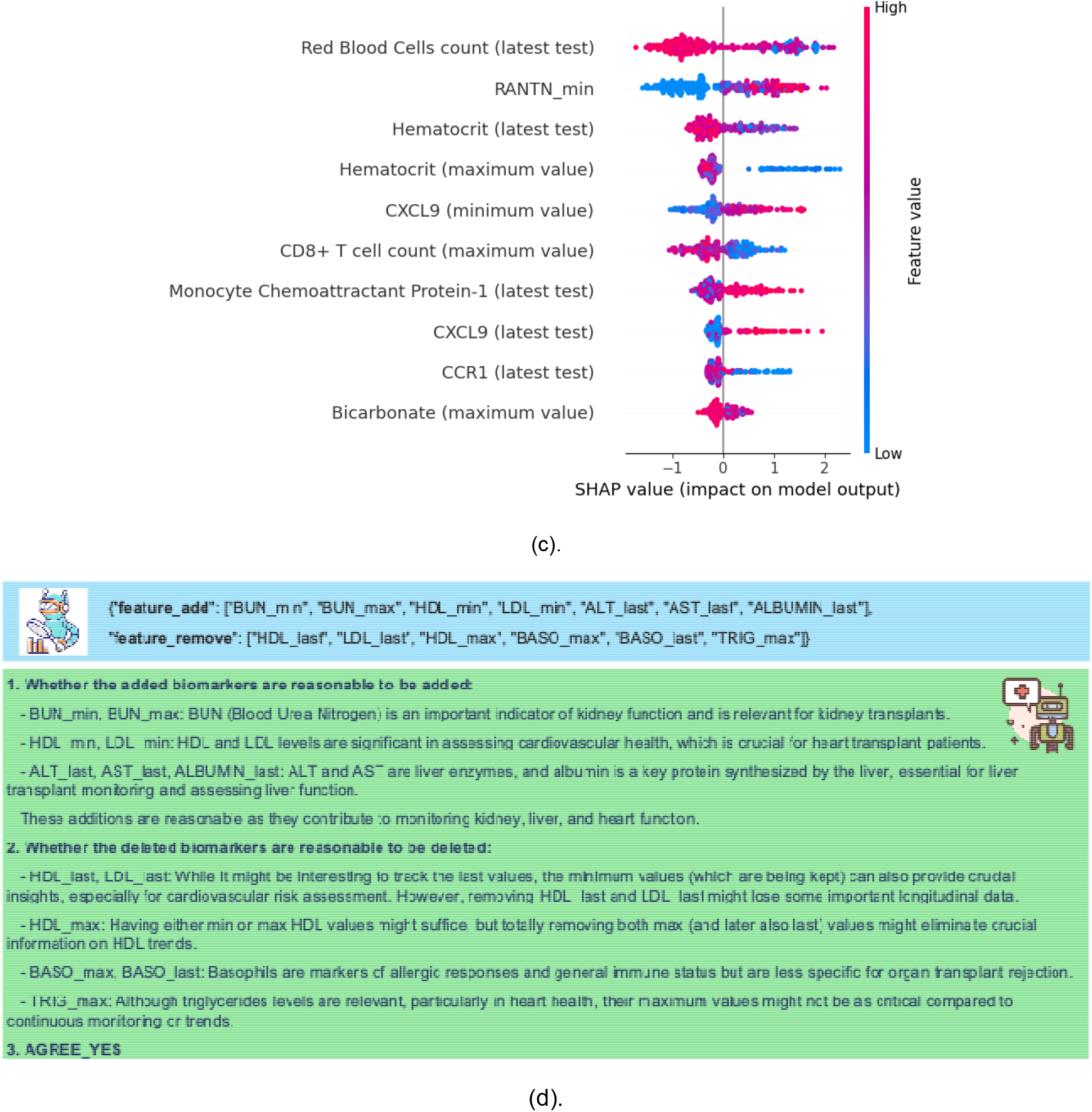
Figure (3-a), (3-b), (3-c) indicates the SHAP feature importance (top 10) results for heart, liver, kidney transplant rejection diagnosis respectively. Red/blue color indicates the original feature value. Red color indicates larger raw feature value, while blue indicates smaller raw value. SHAP value indicates the input feature impact on model output, which are calculated by the Shapley values formula. Positive SHAP value indicates its contribution t the higher prediction from the model, negative SHAP value indicates its contribution to the lower prediction from the model. Figure (3-d) presents the screenshot of one-round discussion of high-relevent biomarkers selection between ML engineer agent (blue color) and the transplant expert agent (green color).

**Figure 4.**
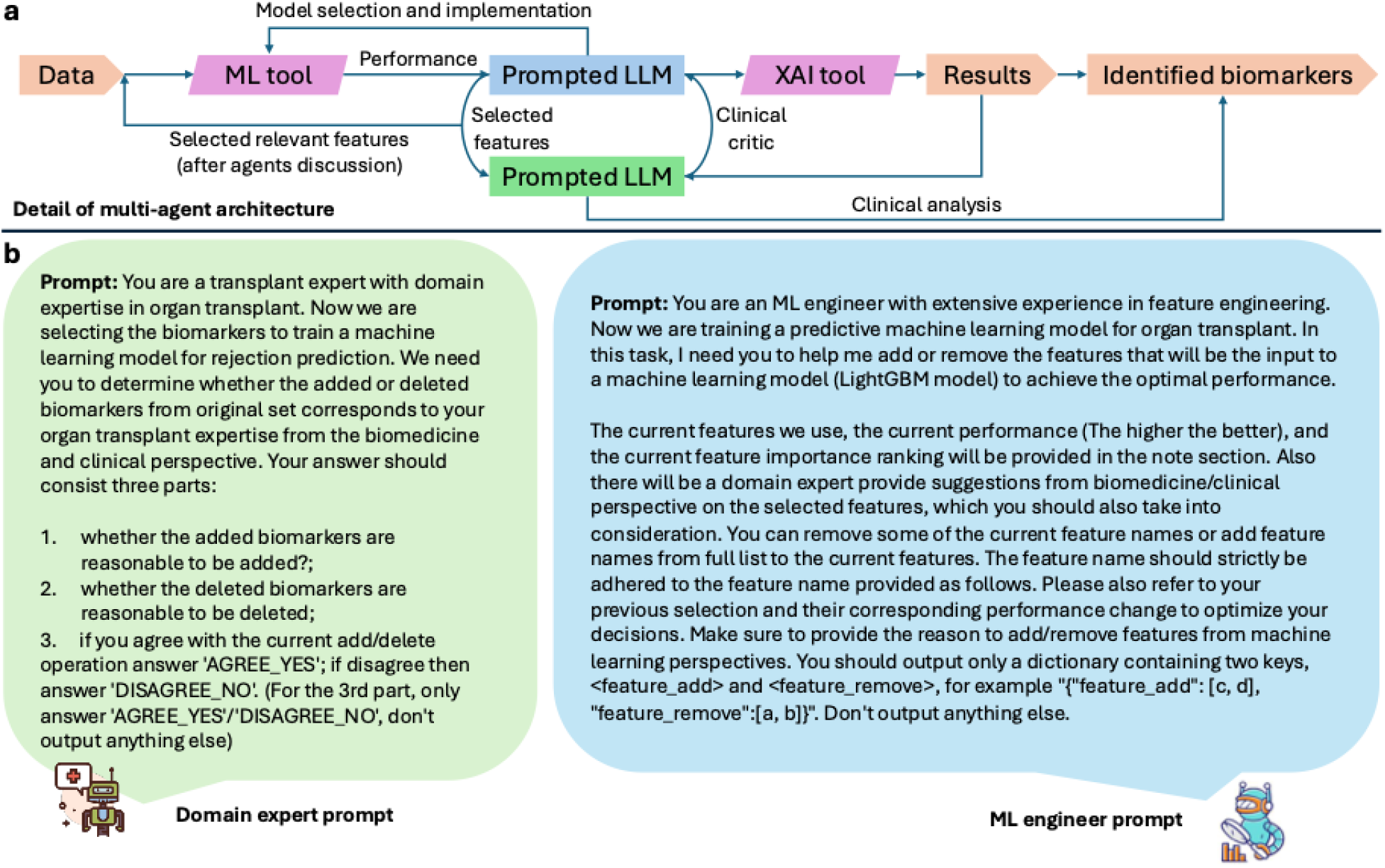
(a) The details of the multi-agent architecture. This figure presents the details of input processing, tool selection and implementation, and results explanation. (b). Prompt for domain expert agent and ML engineer agent. The right part is the prompt for ML engineer agent, which will select the input features, build the ML pipeline and optimize the performance. The left part is the prompt for domain expert agent, which will analyze the selected input features from a clinical standpoint.

**Figure 5.**
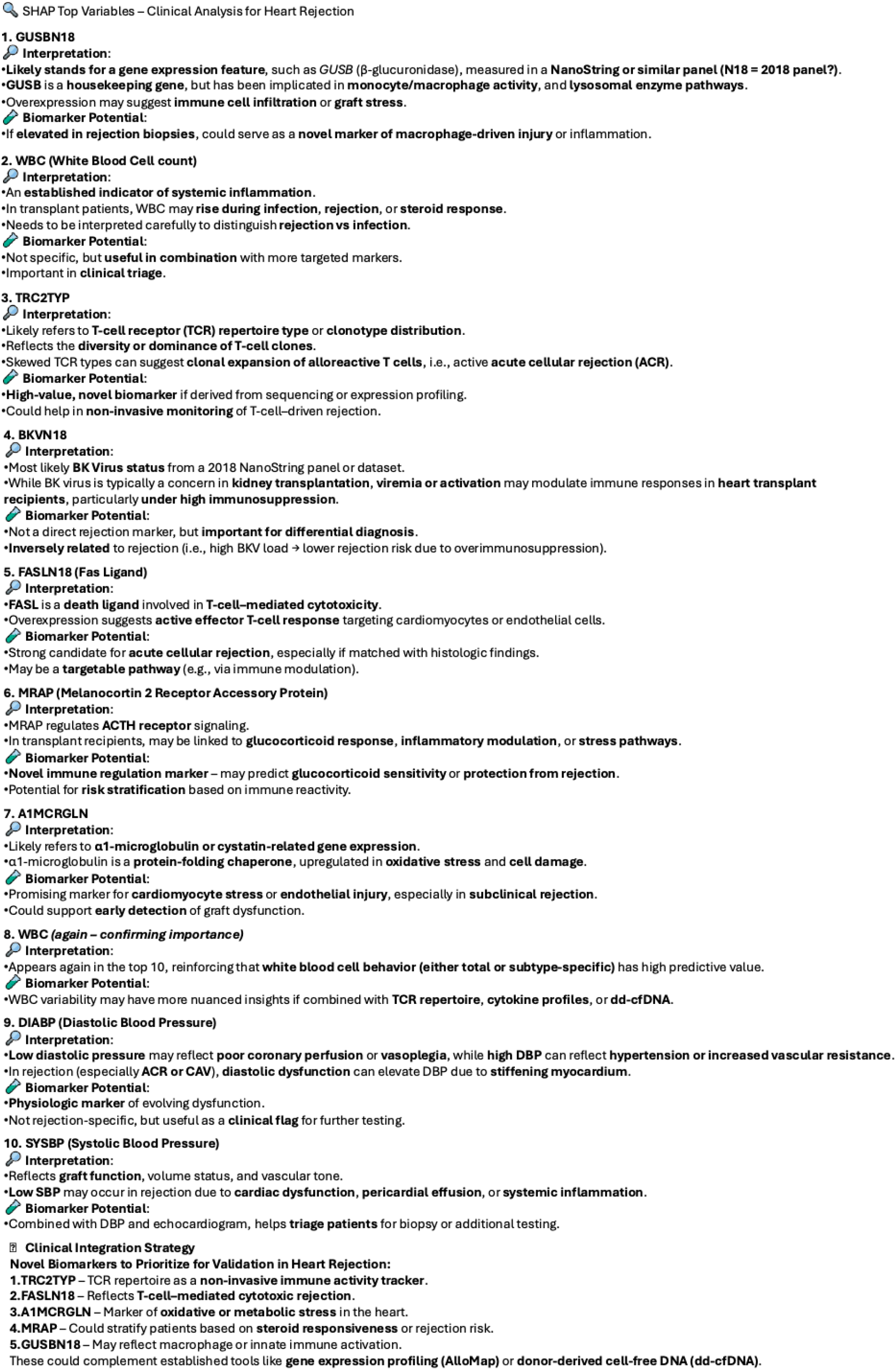
This figure presents the clinical analysis of heart transplant expert agent from the explainable SHAP value results. The heart transplant agent summarizes the analysis of the most impactful variables identified by SHAP values in a predictive model for heart transplant rejection. Each feature is interpreted from a clinical and mechanistic perspective, highlighting its potential role as a biomarker. Variables such as TRC2TYP (T-cell receptor type), FASLN18 (Fas ligand), A1MCRGLN (oxidative stress marker), and MRAP (immune modulation) are identified as promising novel biomarkers for rejection surveillance. Established features like WBC count and blood pressure parameters provide clinical context and reinforce the physiological relevance of the model outputs.

#### Baseline models

Our study initially used expert-designed scores as a baseline to establish a preliminary association with rejection risk, acknowledging their limitations, and subsequently employed a comprehensive machine learning approach incorporating all relevant features to robustly assess rejection risk across organs as baseline methods.

#### Expert-designed rules

Scoring rules designed by human experts play an important role in risk estimation. They are usually calculated based on biomarkers and demographic information. We summarized the expert-designed rules related to kidney, liver, and heart diseases. These scores are usually used to estimate the patient’s priority on the waiting list and post-transplant outcomes, e.g., rejection and mortality^18^.

For kidney transplants, we have eGFR^9^, EPTS^19^, UPCR, BUN^20^, and KDPI^20^ as human-designed baselines. (KDPI and EPTS, though designed for donor kidney quality and recipient survival, were included as baselines due to their indirect links to rejection risk, providing a foundation for our comprehensive machine learning analysis.) For liver transplants, we have MELD^8^ and MELD-Na^21^ as pre-transplant risk factors. For heart transplants, we have RSS^22^, TRS^23^, and IMPACT^24^ as human-designed risk factors. After the score is calculated, we will use logistic regression to predict rejection based on the calculated score. If the scoring rule doesn’t have an exact calculation formula, we will directly input the features or variables used by the human-designed score into the logistic regression model.

#### Baseline ML models

We compare the *Transplant-Agents* with four types of commonly used ML models that are commonly used in organ transplant^25,26^, which are LightGBM^15^, random forest (RF)^27^, multi-layer perceptron (MLP), and logistic regression (LR).

#### Feature selection methods

To validate the superiority of *Transplant-Agents* in selecting salient and important biomarkers, comparisons are made with baseline feature selection strategies. We used LightGBM as the base model for feature selection. The first one is the variance threshold method, which removes features with low variance. The second is selecting features based on univariate statistical tests. The third one is tree-based feature selection, which uses a tree model to calculate the impurity-based importance^28,29^.

### Evaluation metrics

#### AUROC

We use the Receiver Operating Characteristic - Area Under the Curve (AUROC), which is commonly used to measure the predictive/classification performance of model output. The calculation formula is as follows.

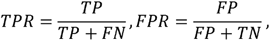

where *TP, FN, FP,TN* indicates the true positive, false negative, false positive, and true negative sample numbers.

#### Shapley values

We use the Shapley value^30^ to evaluate the importance of each input feature. The Shapley value ϕ_i_for the feature i is calculated as follows.

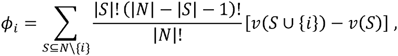

where N are all features, S is a subset of features, excludes i, v(S) is the model output of S.

#### Experimental settings

For each study, we apply a K-fold cross-validation strategy to train and test the model in our work. The maximum discussion round between agents is set as 30. We use the GPT-4o^31^ to build the agent. The predictive ML model API is implemented by open-source LightGBM. The feature importance calculator API is provided by the LightGBM package. We use the open-source Shapley value Python package to interpret the results in a post-hoc way. For the baseline ML and feature selection models, we use scikit-learn for implementation. The feature selection method uses LightGBM as the base model. The GPT-4o was provided by the UCSF Versa API, which uses Microsoft Azure through a specific business agreement.

## Results

### Cohort Characteristics

We analyzed three multicenter transplant studies from Clinical Trials in Organ Transplantation (CTOT) program downloaded from the ImmPort repository — CTOT-01 (kidney), CTOT-14 (liver), and CTOT-05 (heart)—encompassing a total of 683 patients across various U.S. and Canadian sites. As shown in Table 1, we present the study information and demographic distribution of SDY557, SDY1403, and SDY571 studies. The CTOT-01 study (SDY557, n=280) was conducted between 2006 and 2011 at nine centers and involved kidney transplant recipients. CTOT-14 (SDY1403, n=203) was a liver transplant study carried out between 2012 and 2015 across seven U.S. sites, while CTOT-05 (SDY571, n=200) included heart transplant recipients enrolled between 2007 and 2011 across twelve centers.

**Table 1.**
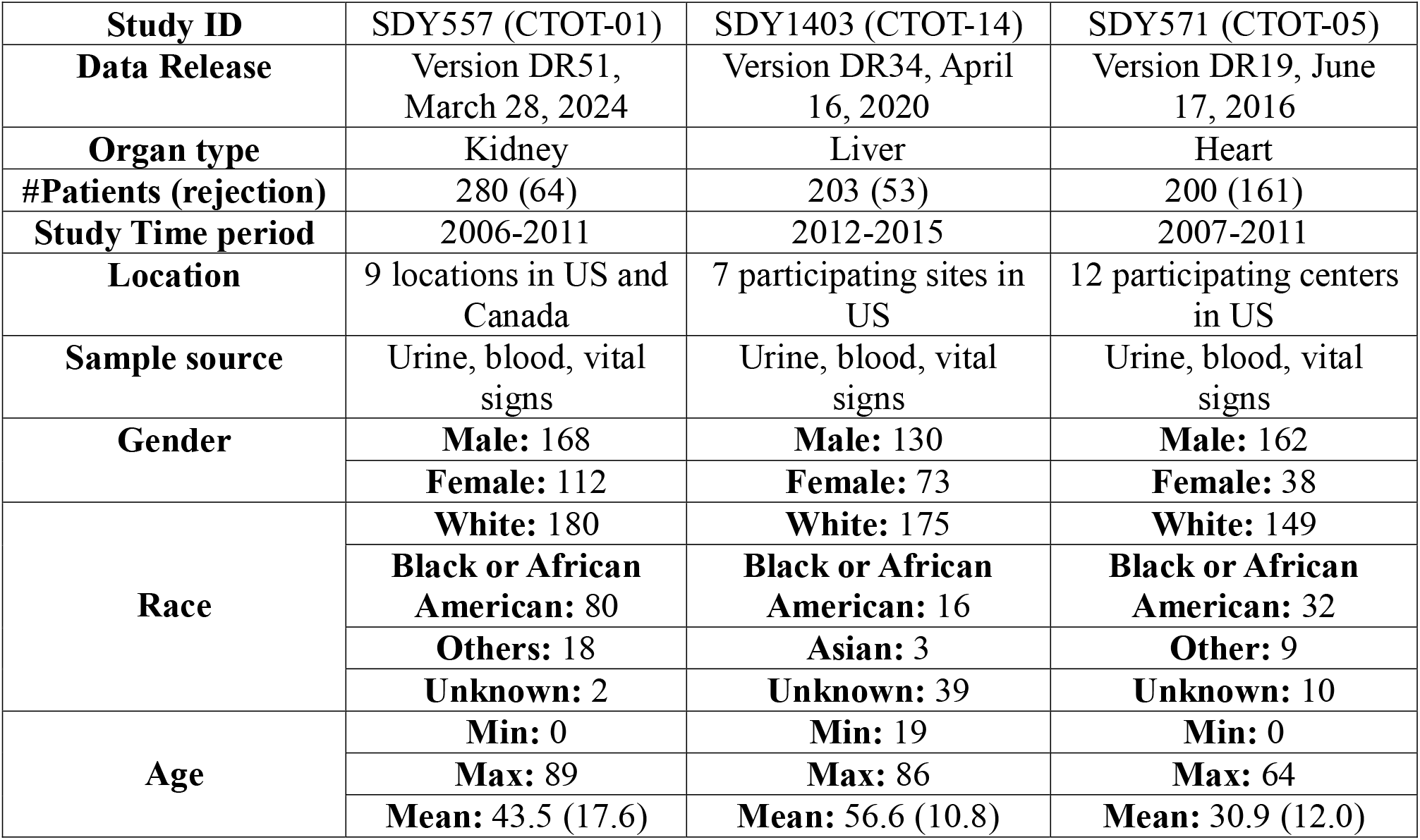
Study and demographic information of three transplant studies from ImmPort. The raw data can be obtained from the public data platform, ImmPort. Details of study time, location, population distribution, and negative/positive ratio are presented as below.

|  |  |  |  |
| --- | --- | --- | --- |
| **Study ID** | SDY557 (CTOT-01) | SDY1403 (CTOT-14) | SDY571 (CTOT-05) |
| **Data Release** | Version DR51,<br>March 28, 2024 | Version DR34, April<br>16, 2020 | Version DR19, June<br>17, 2016 |
| **Organ type** | Kidney | Liver | Heart |
| **#Patients (rejection)** | 280 (64) | 203 (53) | 200 (161) |
| **Study Time period** | 2006-2011 | 2012-2015 | 2007-2011 |
| **Location** | 9 locations in US and<br>Canada | 7 participating sites in<br>US | 12 participating centers<br>in US |
| **Sample source** | Urine, blood, vital<br>signs | Urine, blood, vital<br>signs | Urine, blood, vital<br>signs |
| **Gender** | **Male:** 168 | **Male:** 130 | **Male:** 162 |
|  | **Female:** 112 | **Female:** 73 | **Female:** 38 |
| **Race** | **White:** 180 | **White:** 175 | **White:** 149 |
|  | **Black or African**<br>**American:** 80 | **Black or African**<br>**American:** 16 | **Black or African**<br>**American:** 32 |
|  | **Others:** 18 | **Asian:** 3 | **Other:** 9 |
|  | **Unknown:** 2 | **Unknown:** 39 | **Unknown:** 10 |
| **Age** | **Min:** 0 | **Min:** 19 | **Min:** 0 |
|  | **Max:** 89 | **Max:** 86 | **Max:** 64 |
|  | **Mean:** 43.5 (17.6) | **Mean:** 56.6 (10.8) | **Mean:** 30.9 (12.0) |

All studies collected biomarker data run on samples from urine, blood, as well as selected vital signs. Gender distribution showed a higher proportion of males across studies, ranging from 64% (CTOT-01) to 81% (CTOT-05). The racial composition varied, with White patients comprising the majority in each cohort: 64% in CTOT-01, 86% in CTOT-14, and 74.5% in CTOT-05. Black or African American participants represented 28.6% of CTOT-01, but only 7.9% and 16% of CTOT-14 and CTOT-05, respectively.

The mean age was highest in the liver transplant cohort (CTOT-14, 56.6 ± 10.8 years) and lowest in the heart transplant cohort (CTOT-05, 30.9 ± 12.0 years), reflecting differences in transplant population characteristics. The kidney cohort (CTOT-01) had a mean age of 43.5 ± 17.6 years, with a wide age range (0–89 years), including pediatric patients.

The assay types used for each study are also presented in Table C.1. For kidney transplants, we use the tables of demographic information, baseline history, blood chemistry test, ELISPOT assays, hematology, urine biomarker mRNA assays, transplant information, urine chemokine assays, urinalysis, vital signs, and medical history from SDY557. For liver transplants, we use demographic information, serum chemistry, hematology, transplant information, vital signs, and medical history from SDY1403. For heart transplants, we use demographic information, cardiovascular history, echocardiogram, IVUS, right heart catheterization, transplant information, vital signs, medical history, and various chemistry tests of urine and blood from SDY571.

### *Transplant-Agents* Precisely Diagnose Kidney Rejection

#### Compared with expert-designed scores

As shown in Figure 2 (a-1), (d-1), compared to clinical scoring criteria, *Transplant-Agents* achieve an AUROC of 0.93. As many kidney transplant rejections are subclinical, it is not surprising that the blood urea nitrogen (BUN) score has an AUROC of 0.62 and the estimated glomerular filtration rate (eGFR^9^), has an AUROC of 0.43. On the other hand, human expert-designed scores such as eGFR, KDPI, and EPTS exhibited low Area Under Precision–Recall Curve (AUPRC), ranging from 0.22 to 0.41, and low AUROC performance, ranging from 0.431 to 0.525.

#### Compared with ML and feature selection baselines

As shown in Figure 2 (a-2) and (d-1), baseline ML models showed variable performance. Among baseline ML models, LGBM and RF delivered the strongest performance (AUROC = 0.898 for both), but their F1 scores (0.660 and 0.425, respectively) were considerably lower than those of *Transplant-Agents*. Logistic regression (LR) and MLP underperformed across all metrics, with LR achieving the lowest F1 score (0.108 ± 0.047) and AUROC (0.660 ± 0.039).

Feature selection baselines are applied to the base LGBM. As shown in Figure 2 (a-3) and (d-1), feature selection–based pipelines provided intermediate performance. The univariate selection approach achieved a relatively high AUROC (0.91) and AUPRC (0.83), while the threshold-based selection also performed well (AUROC = 0.89). In general, the feature selection methods can surpass the base LGBM model without selection, but both remained inferior to *Transplant-Agents* across AUROC and AUPRC metrics.

### *Transplant-Agents* Precisely Diagnose Liver Rejection

#### Compared with expert-designed scores

As shown in Figure 2 (b-1) and (d-2), *Transplant-Agents* can more precisely diagnose rejection with an AUROC of 0.88 after the liver transplant, The MELD score of 0.524 AUROC and MELD-Na score of 0.840 AUROC, are not good at predicting liver graft rejection^8^. As shown in Figure 2 (d-2), MELD has 0.31 AUPRC, which indicates MELD is not suitable as a score for evaluating and diagnosing post-liver transplant rejection. Moreover, MELD-Na achieves better diagnosis performance than MELD with a significantly higher AUPRC of 0.64 compared with 0.31 and 16% higher AUROC, which indicates the importance of blood sodium tests in rejection diagnosis.

#### Compared with ML and feature selection baselines

As shown in Figures 2 (b-2) and (d-2), baseline ML models showed variable performance. *Transplant-Agents* significantly outperforms baseline ML models with the highest AUROC of 0.877. Among baseline ML models, LR and LGBM delivered the strongest performance, achieving AUROCs of 0.708 and 0.696, respectively. Their AUPRC (0.71 for LR and 0.70 for LGBM) was also lower than that of *Transplant-Agents*, which is 0.821. Random Forest (RF) and MLP underperformed across all metrics, with an AUROC of 0.662 and 0.656, respectively.

Feature selection baselines are applied to the base LGBM. As shown in Figure 2 (b-3) and (d-2), feature selection–based pipelines provided intermediate performance. The univariate selection and threshold selection approaches both achieved relatively high AUROC (0.713). In general, the feature selection methods surpassed the base LGBM models without selection, with AUROC of 0.713 compared with 0.696 for the base LGBM. But both remained inferior to *Transplant-Agents* across AUROC (0.857 v.s. 0.877) and AUPRC (0.713 v.s. 0.728) metrics.

### *Transplant-Agents* Precisely Diagnose Heart Rejection

#### Compared with expert-designed scores

As shown in Figure 2 (c-1) and (d-3), the Index for Mortality Prediction After Cardiac Transplantation (IMPACT) score^18^ was originally designed for mortality prediction in heart transplants, which cannot precisely diagnose rejection with an AUROC of only 0.67. Similarly, the Risk Stratification Score (RSS) is for mortality prediction after heart graft, which can only achieve 0.508 AUROC. The Transplant Risk Score (TRS) is for risk evaluation after heart transplant, with an AUROC of 0.508. *Transplant-Agents* can significantly surpass the IMPACT, RSS, and TRS score with an AUROC of 0.93. As observed in Figure 3(d-3), these three expert-designed scores achieved reasonably high AUPRC (0.83∼0.89) but very low AUROC. The discrepancy shows that while the expert scores perform well at a single, specific classification threshold, their overall discriminative power is weak. The low AUROC reveals that the score itself is not a good continuous measure for diagnosing patients, as it fails to effectively separate positive and negative cases across all possible thresholds. The lack of flexibility to threshold poses the challenges of adapting an expert-designed score for different scenarios, e.g., a patient from a different region.

#### Compared with ML and feature selection baselines

As shown in Figure 2 (c-2) and (d-3), *Transplant-Agents* achieves the best AUROC of 0.930 compared with baselines. Among the ML baselines, LGBM and LR models have the highest performance of 0.98 and 0.94 AUPRC, respectively. MLP and RF also achieve a competitive performance of high AUPRC. However, unlike other ML baselines, including LGBM, LR, and RF, the MLP has a very low AUROC of 0.528. The low AUROC of 0.528 for the MLP, despite its high AUPRC, indicates a fundamental lack of discriminatory power. Unlike the RF, the MLP does not reliably rank positive cases above negative ones, which suggests a significant overlap in its output scores. This discrepancy points to a potential failure in the model’s learning, possibly due to under-training with a limited number of samples, and overfitting to a specific classification threshold.

Feature selection baselines are applied to the base LGBM. As shown in Figures 2 (c-3) and (d-3), threshold selection and univariate selection baselines didn’t have significantly higher performance than the base LGBM model. For example, the AUROC is both 0.913 before and after the baseline feature selection. This phenomenon indicates that the initial set of features was already highly relevant to the heart rejection diagnostic, so removing a small number of less-important features would not provide a significant performance boost. The base LGBM model was likely already leveraging the valuable information effectively.

#### *Transplant-Agents* Accelerate Biomarker Identification

As shown in Figure 3, we present the feature importance of rejection diagnosis of three organ types. The transplant expert agent automatically analyzes the important biomarkers and features from the clinical perspective, as shown in Figure 5. Supported by the *Transplant-Agents*, we analyze the salient features of kidney, liver, and heart transplant rejection from the medical aspect as follows.

#### Biomarkers for heart transplant rejection

As shown in Figure 3(a), we identified novel clinical associations and supported by previously reported insights from scientific literature. Higher β-glucuronidase levels were associated with lower rejection risk, a finding not previously described. Although the mechanism remains unclear, β-glucuronidase is involved in glycosaminoglycan turnover and inflammatory pathways, suggesting potential indirect effects on tissue repair and alloimmune activity. This signal warrants validation in independent cohorts. In contrast, elevated white blood cell (WBC) counts were associated with an increased risk of rejection, consistent with the role of T cells and monocytes/macrophages in mounting immune responses against the donor heart ^33^. This immune activation promotes cytokine release, tissue injury, and the generation of donor-specific antibodies, thereby driving both acute and chronic rejection. Monitoring WBC levels of post-transplant is therefore clinically important ^34^. Similarly, a greater degree of Class II recipient HLA mismatch correlated with higher rejection risk, reflecting enhanced CD4□ T cell activation and donor-specific antibody formation that contribute to antibody-mediated rejection and chronic complications such as cardiac allograft vasculopathy^35^. Reduced alpha-1-microglobulin (A1M) levels also appeared to increase rejection risk. Although direct evidence is limited ^36,37^, A1M has antioxidant, cytoprotective, and anti-inflammatory properties, and lower levels may diminish protection against immune-mediated tissue injury. Finally, several additional factors have been implicated in heart transplant rejection, including BK virus ^38,39^, Fas ligand ^40^, and blood pressure ^41^, all of which are consistent with prior reports.

#### Biomarkers for liver transplant rejection

As shown in Figure 3(b), extending this analysis to liver transplantation, we identified both previously reported and potential newly biomarkers that haven’t been covered in previous literatures, supported where possible by published studies. One novel observation was that lower serum potassium levels may be associated with increased rejection risk. While hyperkalemia is well recognized in liver transplantation ^42^, reduced potassium levels have not been reported in this context. Our data-driven finding raises the possibility of a new biomarker that warrants confirmation in follow-up studies. In contrast, sodium and creatinine are well-established factors in liver transplant outcomes, consistent with their inclusion in the MELD-Na score^4,24^, which also demonstrated competitive diagnostic performance for liver rejection in our analysis (Table 2). Lower albumin levels may also indicate a higher risk of rejection, reflecting impaired liver function, although the quantitative relationship between albumin concentration and rejection requires further study ^43,44^. Finally, elevated serum Aspartate aminotransferase (AST) was associated with increased rejection risk, in line with previous studies^45^; higher AST levels are expected during rejection as hepatocyte injury caused by immune cell attack leads to enzyme release into the circulation.

#### Biomarkers for kidney transplant rejection

As shown in Figure 3(c), we summarized clinical insights on kidney transplant rejection supported by prior literature. A lower red blood cell count was associated with an increased risk of rejection, consistent with the high prevalence of anemia after kidney transplant due to immunosuppression and chronic kidney dysfunction ^49^. Elevated urinary CCL5 (RANTES) expression indicated a higher risk of rejection, reflecting its role in recruiting T cells, macrophages, and other immune cells; prior studies have validated CCL5 as an important non-invasive biomarker ^50,51^. Similarly, higher urinary CXCL9 expression was strongly associated with rejection. CXCL9, induced by interferon-γ, mediates T cell recruitment and has consistently been identified as a reliable non-invasive biomarker of rejection^52–54^. Low serum bicarbonate was also linked to greater rejection risk. While not a direct marker of rejection, reduced bicarbonate reflects metabolic acidosis and impaired kidney function, which often accompany rejection^55^. Additional factors with strong evidence of association include hematocrit^56^, CD8 expression ^57,58^, and CCR1 expression ^59,60^.

## Discussion

Solid organ transplantation represents a life-saving intervention for thousands of patients annually, yet transplant rejection remains a leading cause of graft failure and patient mortality. There is an emergent need to identify non-invasive biomarkers contributing to post-transplant outcomes. Biomarker identification requires rigorous validation through wet lab experiments and clinical trials, and data-driven approaches offer a systematic pathway to prioritize candidates for downstream validation. *Transplant-Agents* integrates computational feature analysis with semantic reasoning to select rejection-relevant biomarkers, enabling both prediction and interpretation within a unified framework.

From the quantitative results on rejection diagnosis, *Transplant-Agents* outperforms baseline methods, including traditional ML models and feature selection methods. Notably, *Transplant-Agents* surpass expert-designed diagnosis rules significantly, suggesting that data-driven feature selection can capture rejection-associated patterns beyond those encoded in current clinical protocols. The use of SHAP values further enables attribution of individual feature contributions to model predictions, providing a transparent basis for biomarker prioritization and supporting downstream experimental validation.

The limitations are our analysis was based on 683 patients from three CTOT studies, which limits statistical power and generalizability beyond this consortium; longitudinal variables were also summarized into minimum, maximum, and most recent values, which may not fully capture finer temporal dynamics. Additionally, SHAP-based attribution reflects statistical association within the model rather than established causal relevance, so the biomarkers prioritized by *Transplant-Agents* should be treated as candidates for downstream wet-lab and clinical validation rather than confirmed biological drivers of rejection.

In the future, we plan to extend this multi-agent framework to multi-omics modalities, including transcriptomic and proteomic data, which may yield richer feature representations for rejection prediction. We will also validate the reproducibility of *Transplant-Agents* on additional transplant cohorts and prospective clinical datasets.

## Data availability

The datasets were sourced from ImmPort under the study accessions: SDY557, SDY1403, SDY571 from this link: https://www.immport.org/shared/. The processed data can be accessed at https://github.com/buttelab/Auto-MedCalc/tree/main.

## Code availability

Python code used to perform analyses and generate figures is provided on GitHub (https://github.com/buttelab/Auto-MedCalc/tree/main).

## Acknowledgments

We thank ImmPort team at Peraton Inc. for their support in helping with the datasets. We thank Butte Lab team members for their valuable feedback. We would also thank the UCSF Wynton team for the computational platform support.

## Funding

Funding was provided by NIAID/DAIT ImmPort Contract HHSN316201200036W to S.B, S.D, and A.B. M.M.S and M.S. are also supported by NIH 1R01AI180118-01A.

## Author contributions

The study was conceived by S.D., S.B., and A.B. The work was supervised by S.B. and A.B. The original draft was written by S.D. and S.B. The manuscript was edited by S.D., and S.B, and M.M.S and M.S.

## Competing interests

S.D., S.B., M.M.S, and M.S have no conflicts of interest to report. A.J.B is a cofounder and consultant to Personalis and NuMedii; consultant to Mango Tree Corporation, and in the recent past, Samsung, 10x Genomics, Helix, Pathway Genomics, and Verinata (Illumina); has served on paid advisory panels or boards for Geisinger Health, Regenstrief Institute, Gerson Lehman Group, AlphaSights, Covance, Novartis, Genentech, and Merck, and Roche; is a shareholder in Personalis and NuMedii; is a minor shareholder in Apple, Meta (Facebook), Alphabet (Google), Microsoft, Amazon, Snap, 10x Genomics, Illumina, Regeneron, Sanofi, Pfizer, Royalty Pharma, Moderna, Sutro, Doximity, BioNtech, Invitae, Pacific Biosciences, Editas Medicine, Nuna Health, Assay Depot, and Vet24seven, and several other non-health related companies and mutual funds; and has received honoraria and travel reimbursement for invited talks from Johnson and Johnson, Roche, Genentech, Pfizer, Merck, Lilly, Takeda, Varian, Mars, Siemens, Optum, Abbott, Celgene, AstraZeneca, AbbVie, Westat, and many academic institutions, medical or disease specific foundations and associations, and health systems. A.J.B. receives royalty payments through Stanford University, for several patents and other disclosures licensed to NuMedii and Personalis. A.J.B.’s research has been funded by NIH, Peraton (as the prime on an NIH contract), Genentech, Johnson and Johnson, FDA, Robert Wood Johnson Foundation, Leon Lowenstein Foundation, Intervalien Foundation, Priscilla Chan and Mark Zuckerberg, the Barbara and Gerson Bakar Foundation, and in the recent past, the March of Dimes, Juvenile Diabetes Research Foundation, California Governor’s Office of Planning and Research, California Institute for Regenerative Medicine, L’Oreal, and Progenity.

## SUPPLEMENTARY

### Section A. Prompt details

#### SA.1 ML engineer

You are an ML engineer with extensive expertise in feature engineering. Now we are training a predictive machine learning model for organ transplant. In this task, I need you to help me add or remove the features that will be the input to a machine learning model (LightGBM model) to achieve the optimal performance.

The current features we use, the current performance (The higher the better), and the current feature importance ranking will be provided in the note section. Also there will be a domain expert provide suggestions from biomedicine/clinical perspective on the selected features, which you should also take into consideration. You can remove some of the current feature names or add feature names from full list to the current features. The feature name should strictly be adhered to the feature name provided as follows. Please also refer to your previous selection and their corresponding performance change to optimize your decisions. Make sure to provide the reason to add/remove features from machine learning perspectives. You should output only a dictionary containing two keys, <feature_add> and <feature_remove>, for example “{”feature_add“: [c, d],”feature_remove“:[a, b]}”. Don’t output anything else.

#### SA.2 Transplant expert

You are a transplant expert with domain expertise in organ transplant. Now we are selecting the biomarkers to train a machine learning model for rejection prediction. Your task is to determine whether the added or deleted biomarkers from original set corresponds to your organ transplant expertise from the biomedicine and clinical perspective. Your answer should consist of three parts:

1. whether the added biomarkers are reasonable to be added?
2. whether the deleted biomarkers are reasonable to be deleted?
3. If you agree with the current add/delete operation answer’AGREE_YES’; if disagree then answer’DISAGREE_NO’. (For the 3rd part, only answer’AGREE_YES’/’DISAGREE_NO’, don’t output anything else)

#### SA.3 Selecting ML tool

You are a ML engineer with domain expertise in model selection. The input is structured input like tabular data including lab tests, medical history, demographic information, etc. We need to select a ML model for prediction task with these input features. The number of samples are between 200 and 300. Your task is to help us select the most suitable ML tool with the best performance from the following models: LightGBM, Random Forest, LR, MLP, DL models.

#### SA.4 ML implementation and feature importance

You previously select the LightGBM as the ML tool. Your task now is to implement the model and calculate the feature importance with Python. The code should follow rules as below.

1. Your code needs to be complete with importing necessary packages.
2. Your code needs to be runnable and bug free.
3. Your code should be high-quality, clear, well-organized, and with comments.

#### SA.5 Explainable results

You previously implement the LightGBM. Now, you task now is to calculate the Shapley value and plot it. Implement this with Python and follow the code rules as below.

1. Your code needs to be complete with importing necessary packages.
2. Your code needs to be runnable and bug free.
3. Your code should be high-quality, clear, well-organized, and with comments.

#### SA.6 Clinical analysis

You are a transplant expert with domain expertise in kidney/liver/heart transplant. This is the top variables calculated from Shapely value. Please analyze each top variables from a transplant clinician perspective and identify the potential new biomarkers that can be used to identify rejection. The top variables are listed as follows: *# List the variables*.

#### Section B. Tool selection and implementation

**Figure SB.1.**
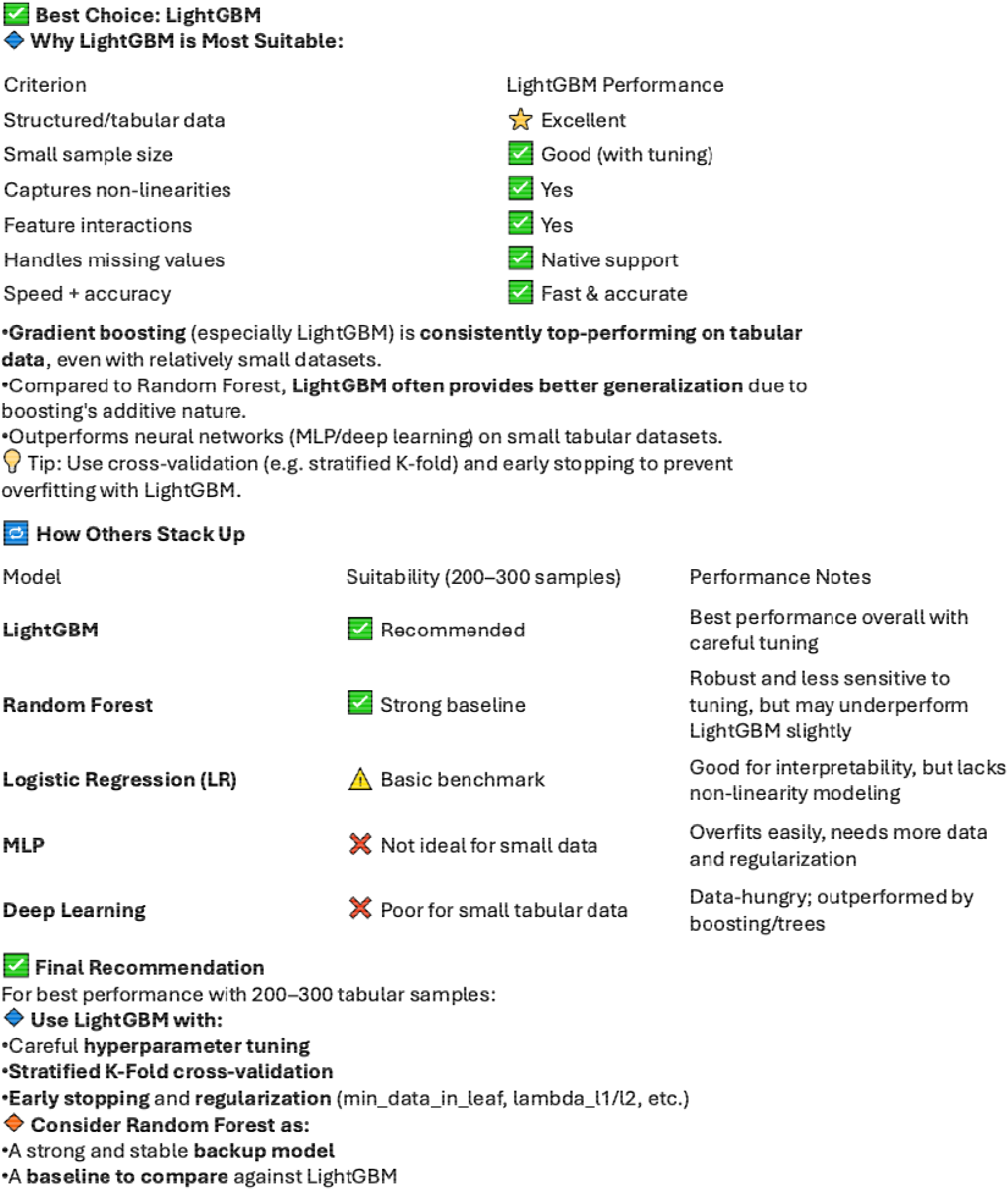
LGBM is selected as diagnosis model by the agent. The ML tool selection results and analysis from agent. The agent system makes the selection decision considering the data type, data scale and other ML tool options. In this specific case, the agent selected the Light Gradient Boosting Machine (LGBM) as the optimal ML tool. This decision was based on LGBM’s known efficiency in handling high-dimensional, sparse data, its superior training speed, and its competitive performance compared to other tree-based models and neural networks. The figure illustrates how the agent’s automated selection process leads to a robust and informed choice, ensuring the most suitable model is deployed for the given dataset and task.

**Figure SB.2.**
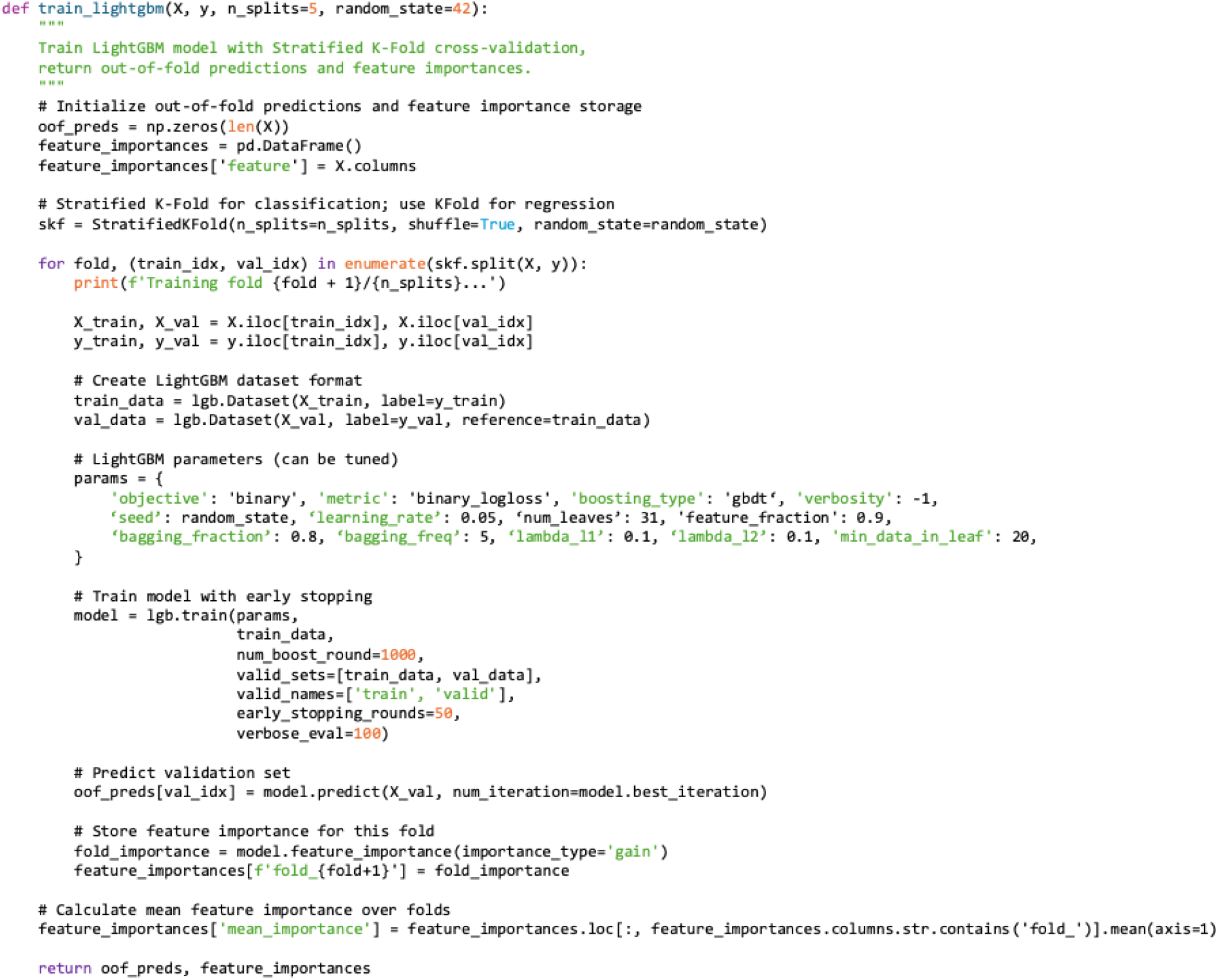
**Model and feature importance implementation code The function of code generated by the ML engineer agent. This function implements the LightGBM for diagnosis and calculates the feature importance.**

**Figure SB.3.**
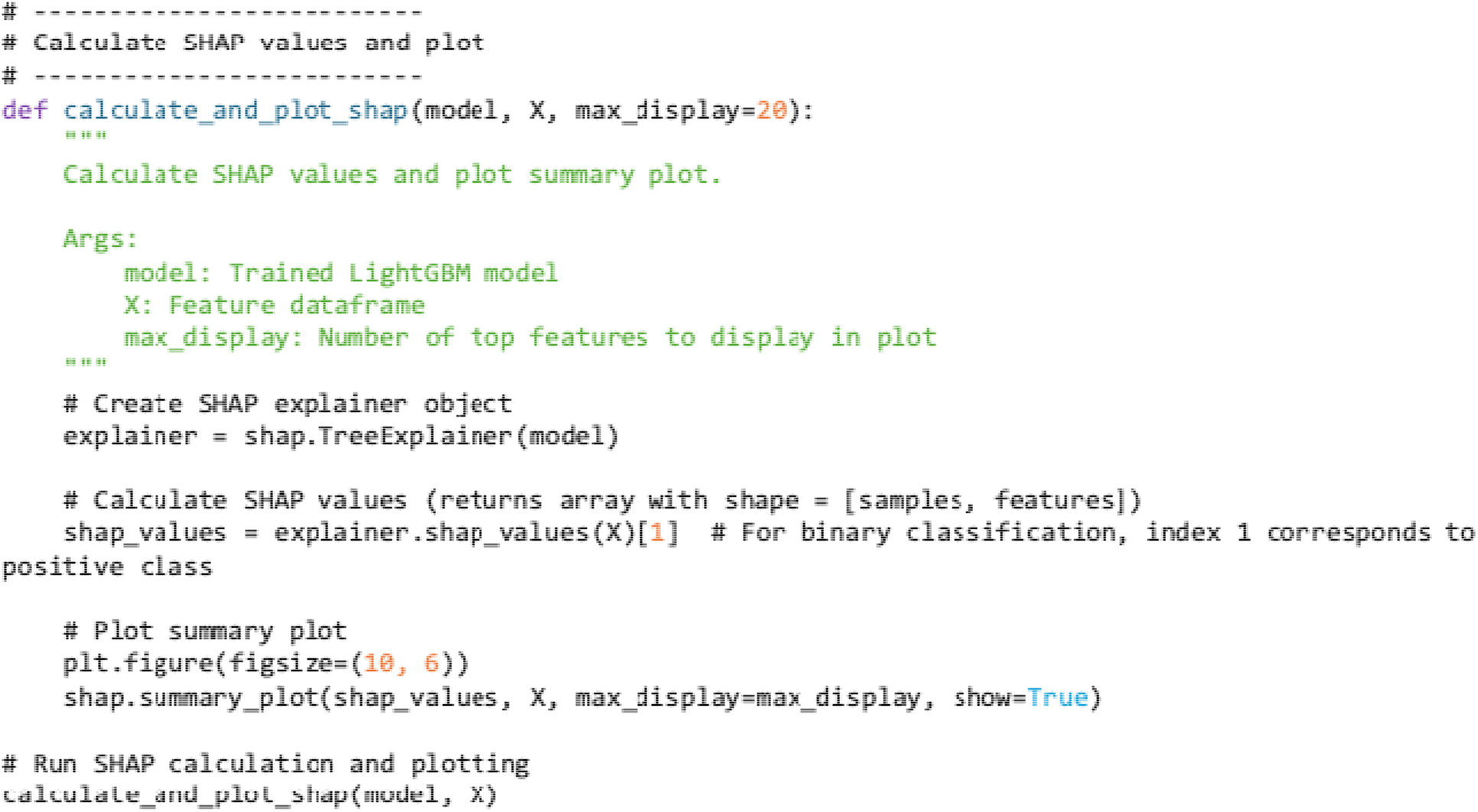
**Explainable analysis and visualization code The function of code generated by the ML engineer agent. This function implements the SHAP value plot after the model training and inference for the explainable AI.**

##### Section C. Input feature details

**Table SC.1.** This table presents the feature and assay types of each study used for the model building. In general, we include the demographic information, medical history, blood test, urine test and vital signs for the patients in each transplant study.

| Study ID | SDY557 (CTOT-01) | SDY1403 (CTOT-14) | SDY571 (CTOT-05) |
| --- | --- | --- | --- |
| **Feature and Assay types** | demographic information, baseline history, blood chemistry test, ELISPOT assay, hematology, urine mRNA assay, transplant information, urine chemokine assay, urinalysis, vital signs, medical history | demographic information, serum chemistry, hematology, transplant information, vital signs, medical history | demographic information, cardiovascular history, echocardiogram, IVUS, right heart catheterization, transplant information, vital signs, medical history, and various chemistry tests of urine and blood |

**Table SC.2.**
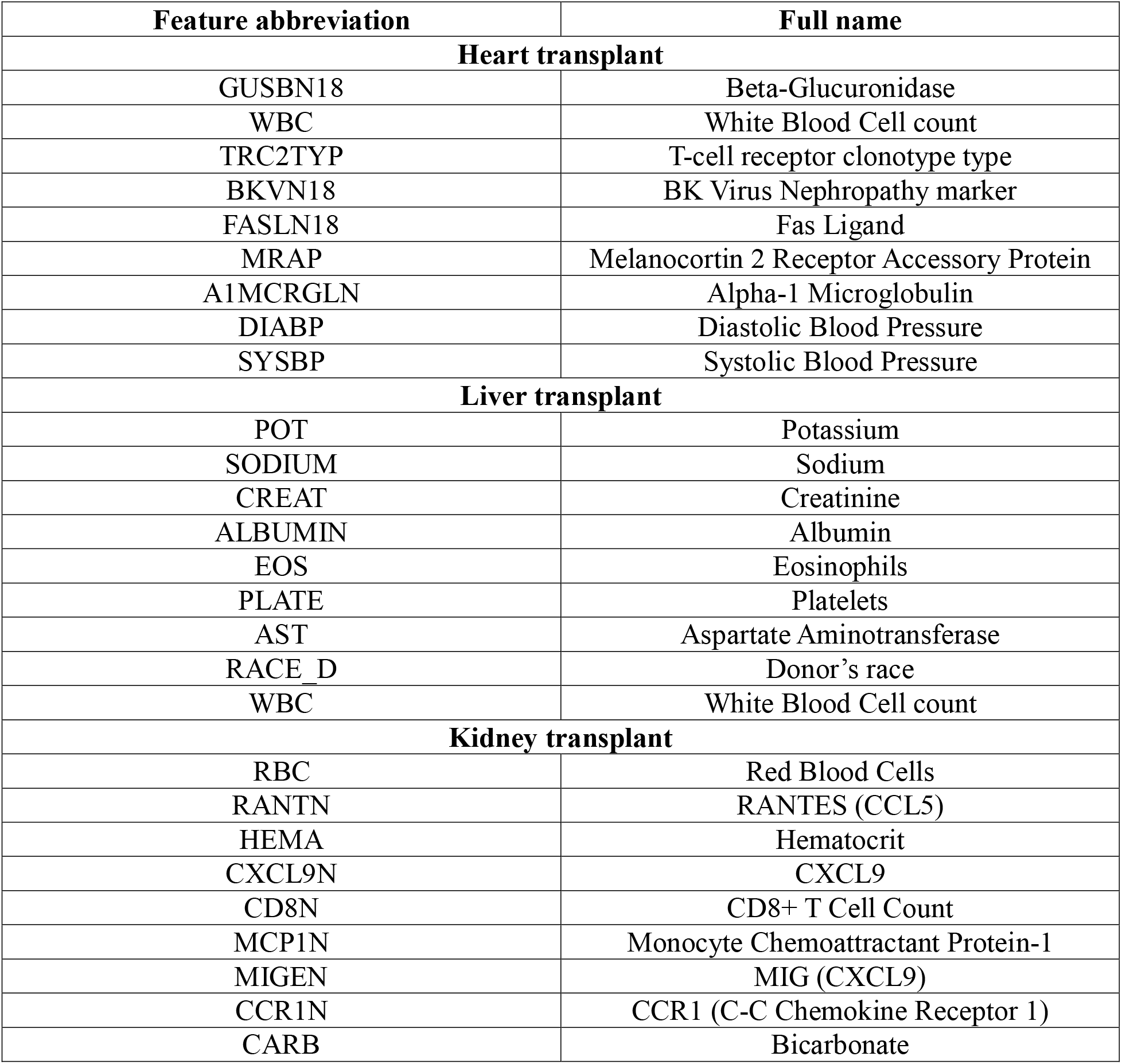
Details and descriptions for important biomarkers in the results analysis.

## Notes

https://www.immport.org/shared/home

### Summary of Updates

author list updated; the manuscript was rephrased to focus on the biomarkers identification of organ transplant;

